# Oxford Nanopore R10.4 long-read sequencing enables near-perfect bacterial genomes from pure cultures and metagenomes without short-read or reference polishing

**DOI:** 10.1101/2021.10.27.466057

**Authors:** Mantas Sereika, Rasmus Hansen Kirkegaard, Søren Michael Karst, Thomas Yssing Michaelsen, Emil Aarre Sørensen, Rasmus Dam Wollenberg, Mads Albertsen

## Abstract

Long-read Oxford Nanopore sequencing has democratized microbial genome sequencing and enables the recovery of highly contiguous microbial genomes from isolates or metagenomes. However, to obtain near-perfect genomes it has been necessary to include short-read polishing to correct insertions and deletions derived from homopolymer regions. Here, we show that Oxford Nanopore R10.4 can be used to generate near-perfect microbial genomes from isolates or metagenomes without shortread or reference polishing.

## MAIN TEXT

Bacteria live in almost every environment on Earth and the global microbial diversity is estimated to entail more than 10^12^ species^1^. To obtain representative genomes, sequencing of pure cultures or genome recovery directly from metagenomes are often employed^2–4^. High-throughput short-read sequencing has for many years been the method of choice^5,6^ but fails to resolve repeat regions larger than the insert size of the library^7^. This is especially problematic in metagenome samples where related species or strains often contain long sequences of near-identical DNA. More recently, long-read sequencing has emerged as the method of choice for both pure culture genomes^8,9^ and metagenomes^10–12^. PacBio HiFi reads combine low error rates with relatively long reads and generate near-perfect microbial genomes from pure cultures or metagenomes^13–15^. Despite very high-quality raw data, the relatively high cost pr. base remains an economic hindrance for many research projects. A widely used alternative is Oxford Nanopore sequencing which offers low-cost long-read data. However, numerous studies have shown that despite vast improvements in raw error rates, assembly consensus sequences still suffer from insertion and deletions in homopolymers that often cause frameshift errors during gene calling^16–18^. A commonly adopted solution has been to include short-read data for post-assembly error correction^12,19^, although it increases the cost and complexity overhead. Another solution has been to apply reference-based polishing to correct frameshift errors^20–22^, but while it provides a practical solution, which allows gene calling, it does not provide true near-perfect genomes.

We first evaluated the ability for Oxford Nanopore R9.4.1 and R10.4 data to obtain near-perfect microbial genomes through sequencing of the ZymoBIOMICS HMW DNA Standard #D6322 (Zymo mock) consisting of 7 bacterial species and 1 fungus. A single PromethION R10.4 flowcell generated 52.3 gbp of data with a modal read accuracy of 99 % (**Figure 1A, Table S1**). In contrast to R9.4.1 data, we do not see any significant improvement in assembly quality for R10.4 by the addition of Illumina polishing (**Figure 1C, Figure S1**). This indicates that near-perfect microbial reference genomes can be obtained from R10.4 data alone at a coverage of approximately 40x. The improvement in assembly accuracy from R9.4.1 to R10.4 is largely due to an improved ability to call homopolymers, as R10.4 is able to correctly call the length of the majority of homopolymers up to a length of 10 (**Figure 1B, Figure S2–3**). In general, a homopolymer length of more than 10 is very rare in bacteria, with an estimate of less than 10 per species on average18.

**Figure 1:**
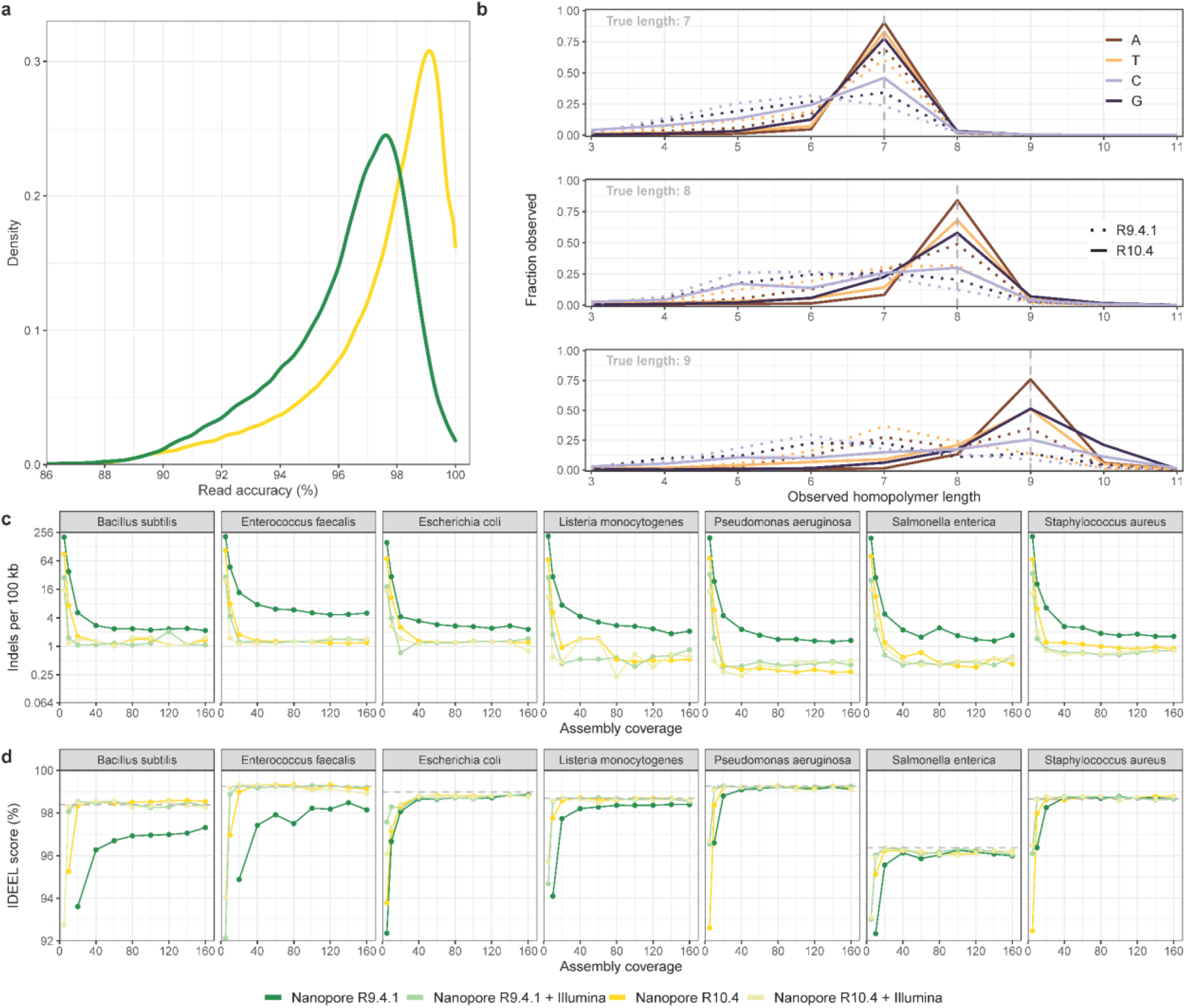
Sequencing and assembly statistics for the Zymo mock. **A)** Observed raw read accuracies measured through read-mapping. **B)** Observed homopolymer length of raw reads compared to the reference genomes (see **Figure S2–3** for a complete overview). **C)** Observed indels of de novo assemblies per 100 kbp at different coverage levels, with and without Illumina polishing. Note that the reference genomes available for the Zymo mock are not identical to the sequenced strains (**Table S3**). **D)** IDEEL^23^ score calculated as the proportion of predicted proteins which are ≥95% the length of their best-matching known protein in a database16. The dotted line represents the IDEEL score for the reference genome.

To assess the performance of state-of-the-art sequencing technologies in recovering near-perfect microbial genomes from metagenomes we sequenced activated sludge from an anaerobic digester using single runs of Illumina MiSeq 2×300 bp, PacBio HiFi, and Oxford Nanopore R9.4.1 and R10.4. Despite being the same sample, direct comparisons are difficult as the additional size selection of the PacBio CCS dataset both increased the read length (**Figure S4**) and altered the relative abundances of the species in the sample (**Figure S5**). Furthermore, Nanopore R9.4.1 produced more than twice the amount of data compared to the other datasets, while the Illumina data featured variations in relative abundances presumably due to GC bias (**Figure S5**). To assist automated contig binning, we performed Illumina sequencing of 9 additional samples from the same anaerobic digester spread over 9 years (**Table S2**) and used the coverage profiles as input for binning using multiple different approaches. Furthermore, to evaluate the impact of micro-diversity on MAG quality, we calculated the polymorphic site rates for each MAG as a simple proxy for the presence of micro-diversity^6^.

After performing automated contig binning it is evident that micro-diversity has a large impact on MAG fragmentation, but that long-read sequencing data results in much less fragmentation of bins at higher amounts of micro-diversity (**Figure S6**). Despite large differences in read length for Nanopore and PacBio CCS data (N50 read length 6 kbp vs. 15 kbp), only small differences in bin fragmentation were observed, as compared to the Illumina-based results (**Table 1, Figures S6**).

**Table 1:** Sequencing and assembly statistics for the anaerobic digester sample using different technologies and approaches. *Costs refer to the expenses encountered at the time of conducting the experiments and may differ for other research groups.

|  | Illumina<br>MiSeq | R9.4.1 /<br>+Illumina | R10.4 /<br>+Illumina | PacBio HiFi |
| --- | --- | --- | --- | --- |
| Total Yield (Gbp) | 13 | 35 | 14 | 15 |
| Read N50 (kbp) | 0.3 | 5.9 | 5.6 | 15.4 |
| Observed modal read accuracy (%) | 100 | 96.76 | 98.21 | 99.86 |
| Assembly size (Mbp) | 409 | 754 | 379 | 606 |
| Contigs (> 1kbp) | 145,976 | 24,680 | 21,585 | 8,989 |
| Circular contigs (> 0.5 Mbp) | 0 | 7 | 3 | 9 |
| Contig N50 (kbp) | 3.5 | 79.9 | 40.1 | 172.5 |
| Reads mapped to contigs (%) | 88.1 | 93.5 | 95.4 | 95.2 |
| HQ MAGs | 8 | 64/86 | 34/36 | 74 |
| MQ MAGs | 83 | 114/95 | 65/67 | 72 |
| Contigs pr. HQ MAG (median) | 184 | 15/16 | 21/21 | 9 |
| Mapped reads in HQ MAGs (%) | 16 | 46/49 | 39/40 | 48 |
| Costs (\$)* | 1,200 | 811/2,011 | 811/2,011 | 4,420 |
| Cost per HQ MAG (\$) | 150 | 13/23 | 24/56 | 60 |

All long-read methods produce high numbers of high-quality (HQ) MAGs, which capture 39-49% of all reads (**Table 1**). Nanopore R9.4.1 is able to produce HQ MAGs as a standalone technology, but Illumina polishing increases the number of HQ MAGs from 64 to 86. For Nanopore R10.4, Illumina polishing increases the number of HQ MAGs from 34 to 36. Using the IDEEL test (**Figure 2**), it can be seen that Illumina polishing results in minor improvements for Nanopore R10.4 above a coverage of 40, and that the Nanopore R10.4 is in the same IDEEL range as PacBio HiFi MAGs. As with sequencing of the Zymo mock, the difference from R9.4.1 to R10.4 is largely due to significantly better accuracy in homopolymers for lengths up to 10 (**Figure S7**).

**Figure 2:**
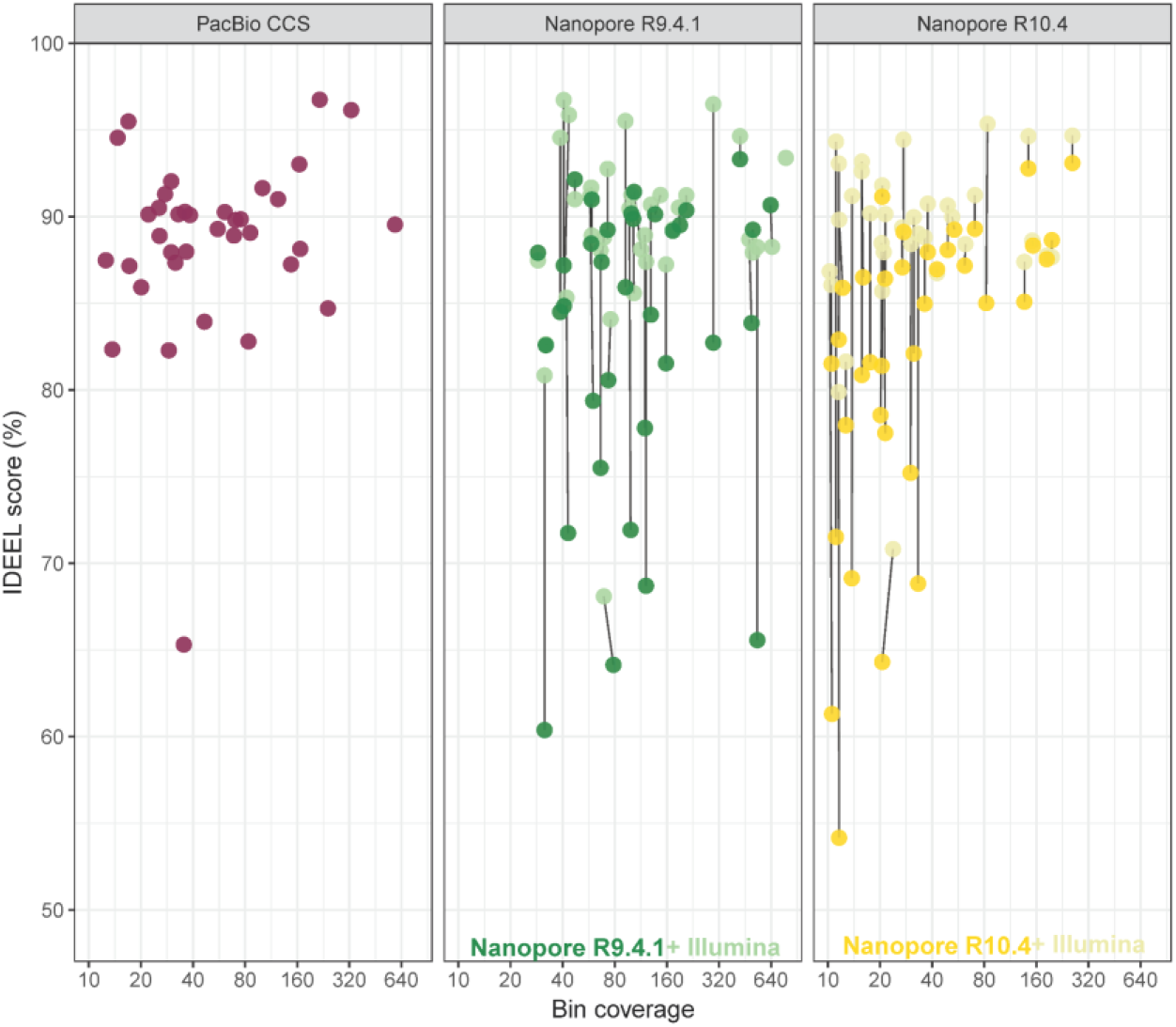
IDEEL score vs. coverage for metagenome bins from the anaerobic digester sample. The Nanopore bins are shown with and without Illumina polishing connected by a line.

Since its introduction as an early access program in 2014 Oxford Nanopore sequencing technology has democratized sequencing and enabled every laboratory and classroom to engage in microbial genome sequencing. However, for the generation of high-quality genomes, additional short-read polishing has been essential, as indels in homopolymer regions cause fragmented gene calls. The additional sequencing requirements have been one of the barriers to widespread uptake. Here we show that Oxford Nanopore R10.4 enables the generation of near-perfect microbial genomes from pure cultures or metagenomes at coverages of 40x without short-read polishing. While homopolymers of 10 or more bases will likely still be problematic, they constitute a minor part of microbial genomes.

For genome-recovery from metagenomes, low-coverage bins (<40X) do need Illumina polishing to attain quality comparable to PacBio HiFi. Hence, in some cases, the most economic option could be Nanopore R9.4.1 supplemented with short-read sequencing, as the throughput is currently at least 2 times higher on R9.4.1 compared to R10.4 and no difference is seen between the methods after Illumina short-read polishing.

## Data availability

Anaerobic digester sequencing data are available at the ENA with bio project ID PRJEB48021, while the Zymo mock community sequencing data is available at PRJEB48692. The code and datasets used to generate the figures and supplementary material are available at https://github.com/Serka-M/Digester-MultiSequencing.

## Acknowledgments

We would like to acknowledge the plant operators at Fredericia wastewater treatment plant for supplying the sample material. The study was funded by research grants from VILLUM FONDEN (15510) and the Poul Due Jensen Foundation (Microflora Danica).

## Author contributions

MS and RHK performed DNA extraction, and sequencing of the anaerobic digester and selected Zymo mock samples. RWO prepared and sequenced the Zymo mock using R9.4.1 and Illumina. MS, RHK, and MA wrote the first draft of the manuscript. SMK, TYM, RWO, and EAS contributed to experiment design, result interpretation, and writing of the manuscript. All authors reviewed the manuscript.

## Materials and methods

### Sampling

Sludge biomass was sampled from the anaerobic digester at Fredericia wastewater treatment plant (Latitude 55.552219, Longitude 9.722003) at multiple time points and stored as frozen 2 mL aliquots at −20°C. For the Zymo sample, the ZymoBIOMICS HMW DNA Standard #D6322 (Zymo Research, USA) was used.

### DNA extraction

DNA was extracted from the anaerobic digester sludge using DNeasy PowerSoil Kit (QIAGEN, Germany) following the manufacturer’s protocol. The extracted DNA was then size selected using the SRE XS (Circulomics, USA), according to the manufacturer’s instructions.

### DNA QC

DNA concentrations were determined using Qubit dsDNA HS kit and measured with a Qubit 3.0 fluorimeter (Thermo Fisher, USA). DNA size distribution was determined using an Agilent 2200 Tapestation system with genomic screentapes (Agilent Technologies, USA). DNA purity was determined using a NanoDrop One Spectrophotometer (Thermo Fisher, USA).

### Oxford Nanopore DNA sequencing

Library preparation was carried out using the ligation sequencing kits (Oxford Nanopore Technologies, UK) SQK-LSK109 and SQK-LSK112 for sequencing on R.9.4.1 and the R.10.4 flowcells, respectively. Anaerobic digester and Zymo R.9.4.1 datasets were generated on a MinION Mk1B (Oxford Nanopore Technologies, UK) device, while Zymo R10.4 dataset was produced on a PromethION and digester R10.4 read sequences were generated on a GridION.

### Illumina DNA sequencing

The anaerobic digester Illumina libraries were prepared using the Nextera DNA library preparation kit (Illumina, USA), while the Zymo Mock sample was prepared with NEB Next Ultra II DNA library prep kit for Illumina (New England Biolabs, USA) following the manufacturer’s protocols and sequenced using the Illumina MiSeq platform.

### PacBio HiFi

A size-selected DNA sample was sent to the DNA Sequencing Center at Brigham Young University, USA. The DNA sample was fragmented with Megaruptor (Diagenode, Belgium) to 15 kb and size-selected using the Blue Pippin (Sage Science, USA) and prepared for sequencing using SMRTbell Express Template Preparation Kit 1.0 (PacBio, USA) according to manufacturers’ instructions. Sequencing was performed on the Sequel II system (PacBio, USA) using the Sequel II Sequencing Kit 1.0 (PacBio, USA) with the Sequel II SMRT Cell 8M (PacBio, USA) for a 30 hour data collection time.

### Read processing

Illumina reads were trimmed for adapters using Cutadapt v. 1.16^24^. The generated raw Nanopore data was basecalled in super-accurate mode with using Guppy v. 5.0.16 (https://community.nanoporetech.com/downloads) with dna_r9.4.1_450bps_sup.cfg model for R9.4.1 and dna_r10.4_e8.1_sup.cfg model for R10.4 chemistry. Concatemers in R10.4 data were split by using “split_on_adapter” command (5 iterations) of duplex-tools v. 0.2.5 (https://github.com/nanoporetech/duplex-tools). Adapters for Nanopore reads were removed using Porechop v. 0.2.3^25^ and reads with Phred quality scores below 7 and 10 for R9.4.1 and R10.4 reads, respectively, were removed using NanoFilt v. 2.6.0^26^. The CCS tool v. 6.0.0 (https://ccs.how/) was utilized with the sub-read data from PacBio CCS to produce HiFi reads. Read statistics were acquired via NanoPlot v. 1.24.0^26^. Zymo read datasets were subsampled to custom coverage profiles using Rasusa v. 0.3.0 (https://github.com/mbhall88/rasusa). Counterr v. 0.1 (https://github.com/dayzerodx/counterr) was used to assess homopolymer calling in reads.

### Read assembly and binning

Long reads were assembled using Flye v. 2.9-b1768^13,27^ with the “--meta” setting enabled and the “--nano-hq” option for assembling Nanopore reads, whereas “--pacbio-hifi” and “-- min-overlap 7500 --read-error 0.01” options were used for assembling PacBio CCS reads, as it resulted in more HQ MAGs than using the default settings. Polishing tools for Nanopore-based assemblies: Minimap2 v. 2.17^28^, Racon v. 1.3.3 (used thrice)^29^, and Medaka v. 1.4.4 (used twice, https://github.com/nanoporetech/medaka). The trimmed Illumina reads were assembled using Megahit v. 1.1.4^30^.

Automated binning was carried out using MetaBAT2 v. 2.12.1^31^, with “-s 500000” settings, MaxBin2 v. 2.2.7^32^ and Vamb v. 3.0.2^33^ with “-o C --minfasta 500000” settings. Contig coverage profiles from different sequencer data as well as 9 additional time-series Illumina datasets of the same anaerobic digester were used for generating the bins. The binning output of different tools was then integrated and refined using DAS Tool v. 1.1.2^34^. CoverM v. 0.6.1 (https://github.com/wwood/CoverM) was applied to calculate the bin coverage (“-m mean” settings) and relative abundance (“-m relative_abundance”) values.

### Assembly processing

The completeness and contamination of the genome bins were estimated using CheckM v. 1.1.2^35^. The bins were classified using GDTB-Tk v. 1.5.0^36^, R202 database. Protein sequences were predicted using Prodigal v. 2.6.3^37^ with “p meta” setting, while rRNA genes were predicted using Barrnap v. 0.9 (https://github.com/tseemann/barrnap) and tRNAscan-SE v. 2.0.5^38^ was used for tRNA predictions. Bin quality was determined following the Genomic Standards Consortium guidelines, wherein a MAG of high quality featured genome completeness of more than 90 %, less than 5 % contamination, at least 18 distinct tRNA genes and the 5S, 16S, 23S rRNA genes occurring at least once^39^. MAGS with completeness above 50 % and contamination below 10 % were classified as medium quality, while low quality MAGs featured completeness below 50 % and contamination below 10 %. MAGs with contamination estimates higher than 10 % were classified as contaminated.

Illumina reads were mapped to the assemblies using Bowtie2 v. 2.4.2^40^ with the “--very-sensitive-local” setting. The mapping was converted to BAM and sorted using SAMtools v. 1.9^41^. Single nucleotide polymorphism rate was then calculated using CMseq v. 1.0.3^6^ from the mapping using poly.py script with “--mincov 10 --minqual 30” settings.

Bins were clustered using dRep v. 2.6.2^42^ with “-comp 50 -con 10 -sa 0.95” settings. Only the bins that featured higher coverage than 10 in their respective sequencing platform and a higher Illumina read coverage than 5 for bins from the hybrid approach were included in downstream analysis. For IDEEL test^17,23^, the predicted protein sequences from clustered bins and Zymo assemblies were searched against the UniProt TrEMBL^43^ database (release 2021_01) using Diamond v. 2.0.6^44^. Query matches, which were not present in all datasets, were omitted to reduce noise. The IDEEL scores were assigned as described by^16^.

QUAST v. 4.6.3^45^ was applied on the Zymo assemblies and the clustered bins with less than 0.5 % SNP rate to acquire mismatch and indels metrics. Cases with Quast parameters “Genome Fraction” of less than 75 % and “Unaligned length” of more than 250 kb were omitted to reduce noise. For homopolymer analysis, the clustered bins were mapped to each other using “asm5” mode of Minimap2 and Counterr was used on the mapping files to get homopolymer calling errors. For QUAST and Counterr, PacBio CCS bins were used as reference sequences. FastANI v. 1.33^46^ was used to calculate identity scores between Zymo assemblies and the Zymo reference sequences. The Zymo mock reference genome sequences were obtained from a link in the accompanying instruction manual to the ZymoBIOMICS HMW DNA Standard Catalog No. D6332 at https://s3.amazonaws.com/zymo-files/BioPool/D6322.refseq.zip.

## Supplementary information

**Figure S1:**
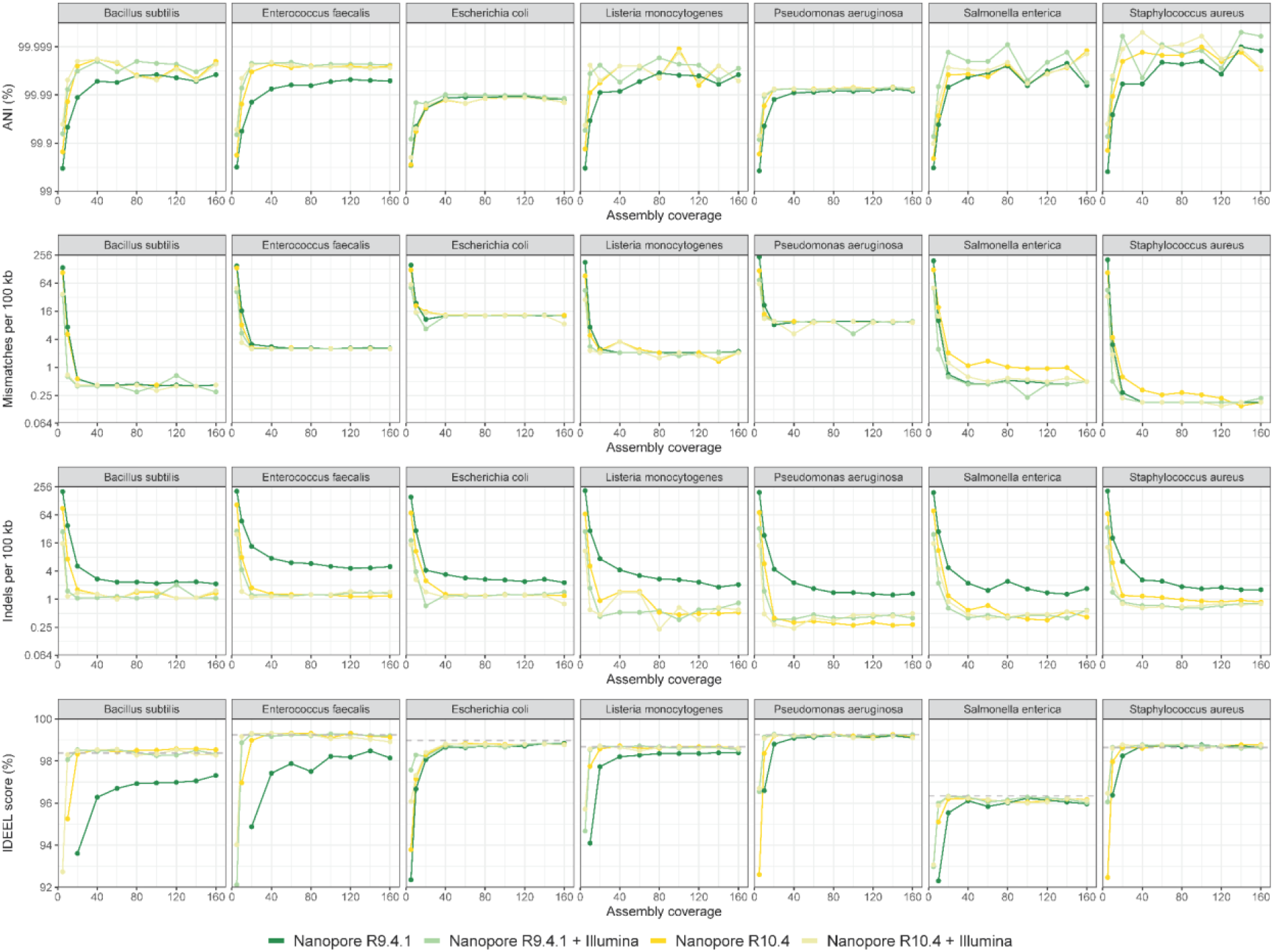
Assembly metrics for the ZYMO Mock HMW DNA.

**Figure S2:**
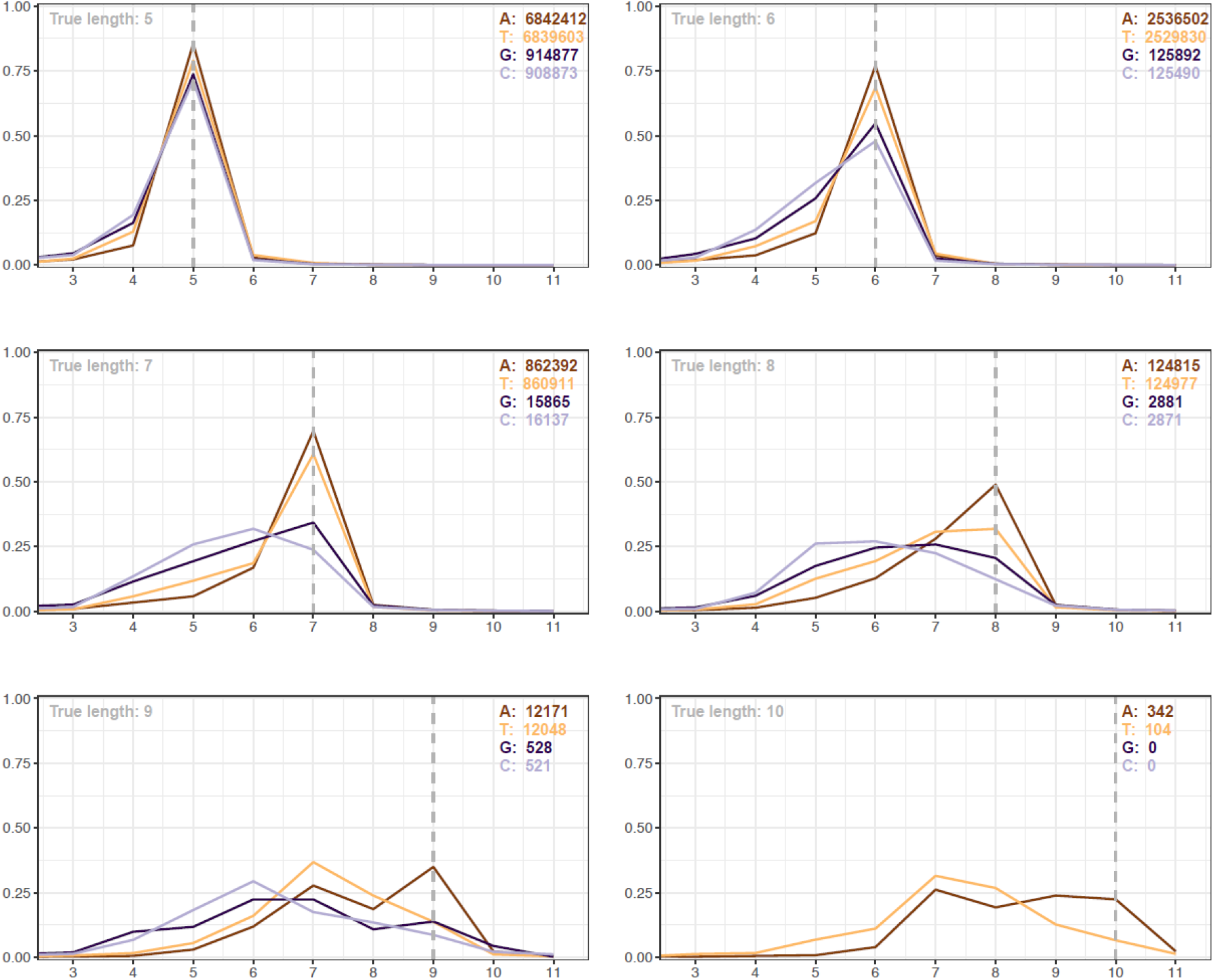
Counterr homopolymer plot for Nanopore R9.4.1 read data of the Zymo mock. Reads for each Zymo mock species, subsetted to a coverage of 160 were used for the analysis.

**Figure S3:**
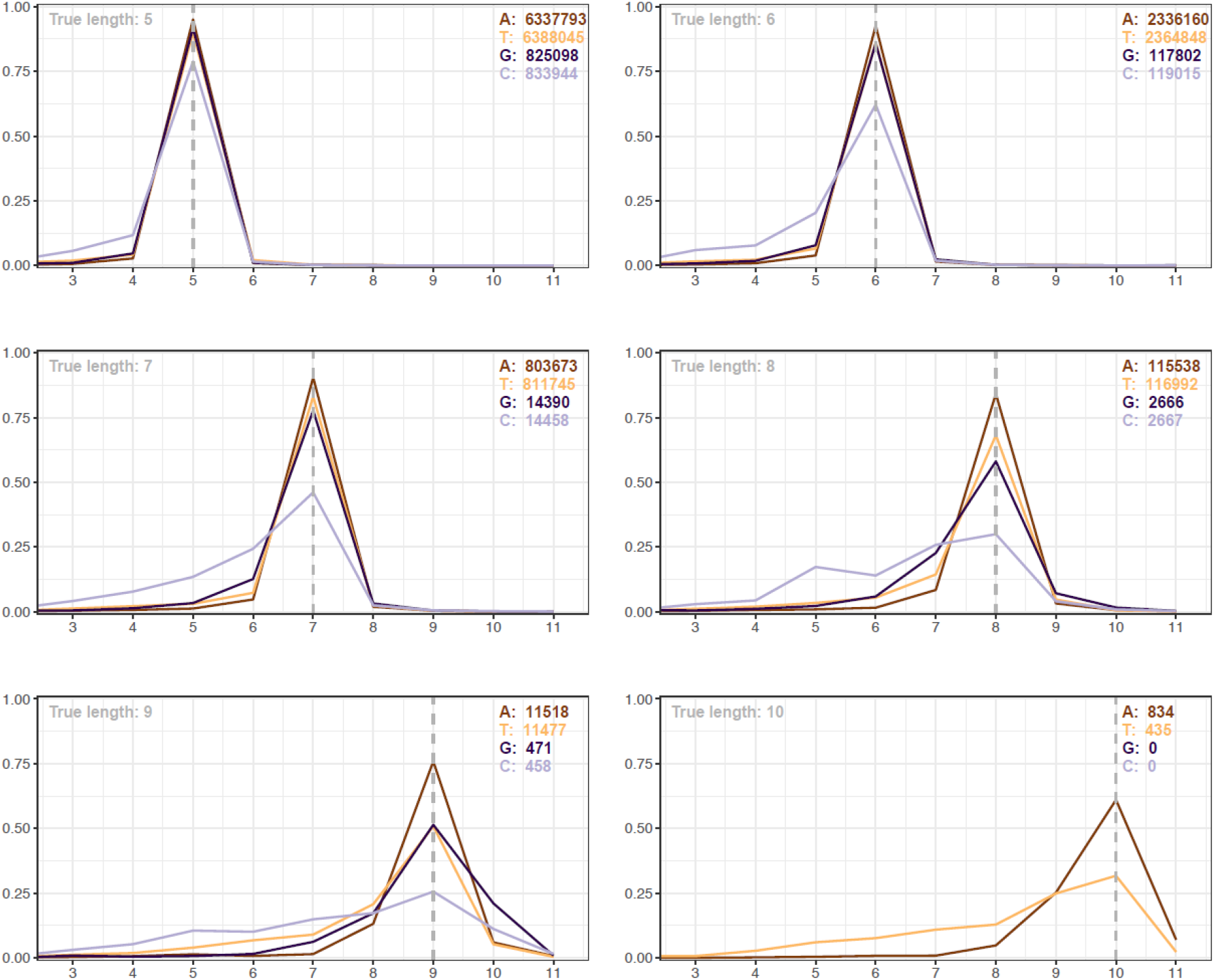
Counterr homopolymer plot for Nanopore R10.4 read data of the Zymo mock. Reads for each Zymo mock species, subsetted to a coverage of 160 were used for the analysis.

**Figure S4:**
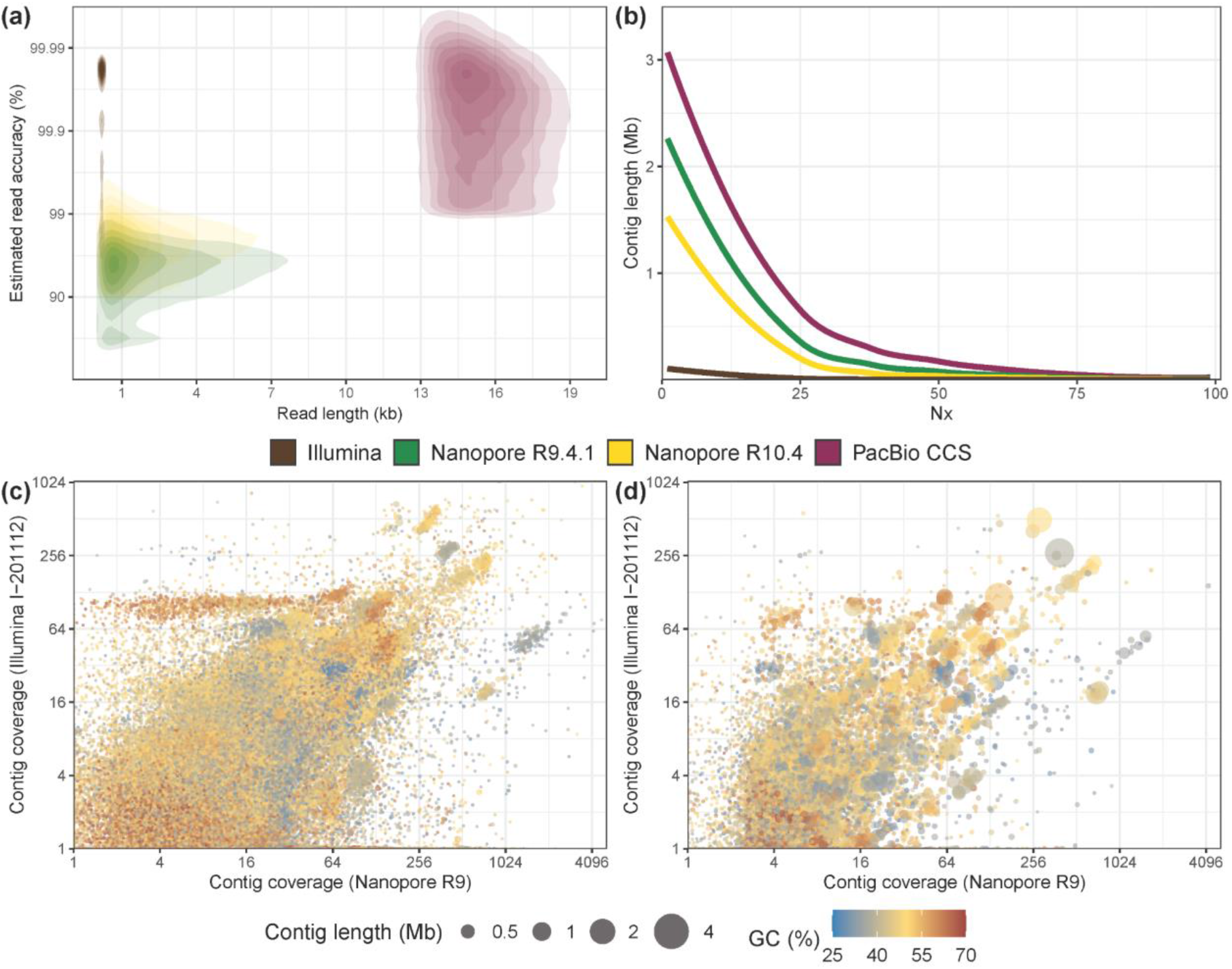
Sequencing and assembly overview for the anaerobic digester sample. **A)** Estimated read accuracy (from Q-scores) versus read length. Note that the PacBio HiFi sample underwent additional size selection prior to sequencing. **B)** Nx plot of the assemblies produced from different sequencing technologies. **C)** Differential coverage plot of the Illumina assembly. **D)** Differential coverage plot of the Nanopore R9.4.1 assembly.

**Figure S5:**
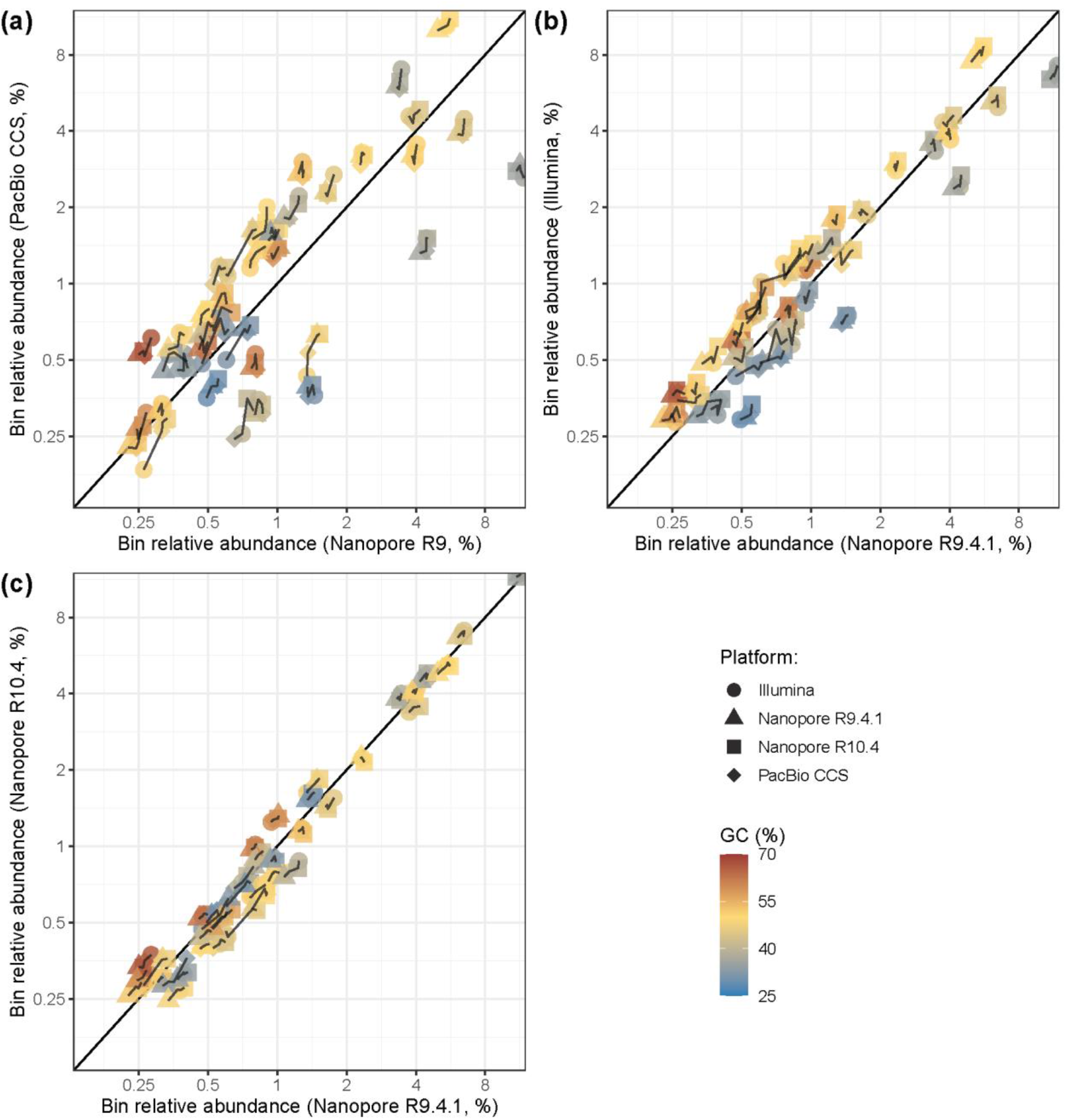
Comparison of bin relative abundances between different sequencing platforms. Relative abundance values (log-scaled) are presented between the Nanopore R9 data and **a)** PacBio CCS, **b)** Illumina, **c)** Nanopore R10. Only the bins that were clustered together between different platforms are presented in the plots and are interlinked.

**Figure S6:**
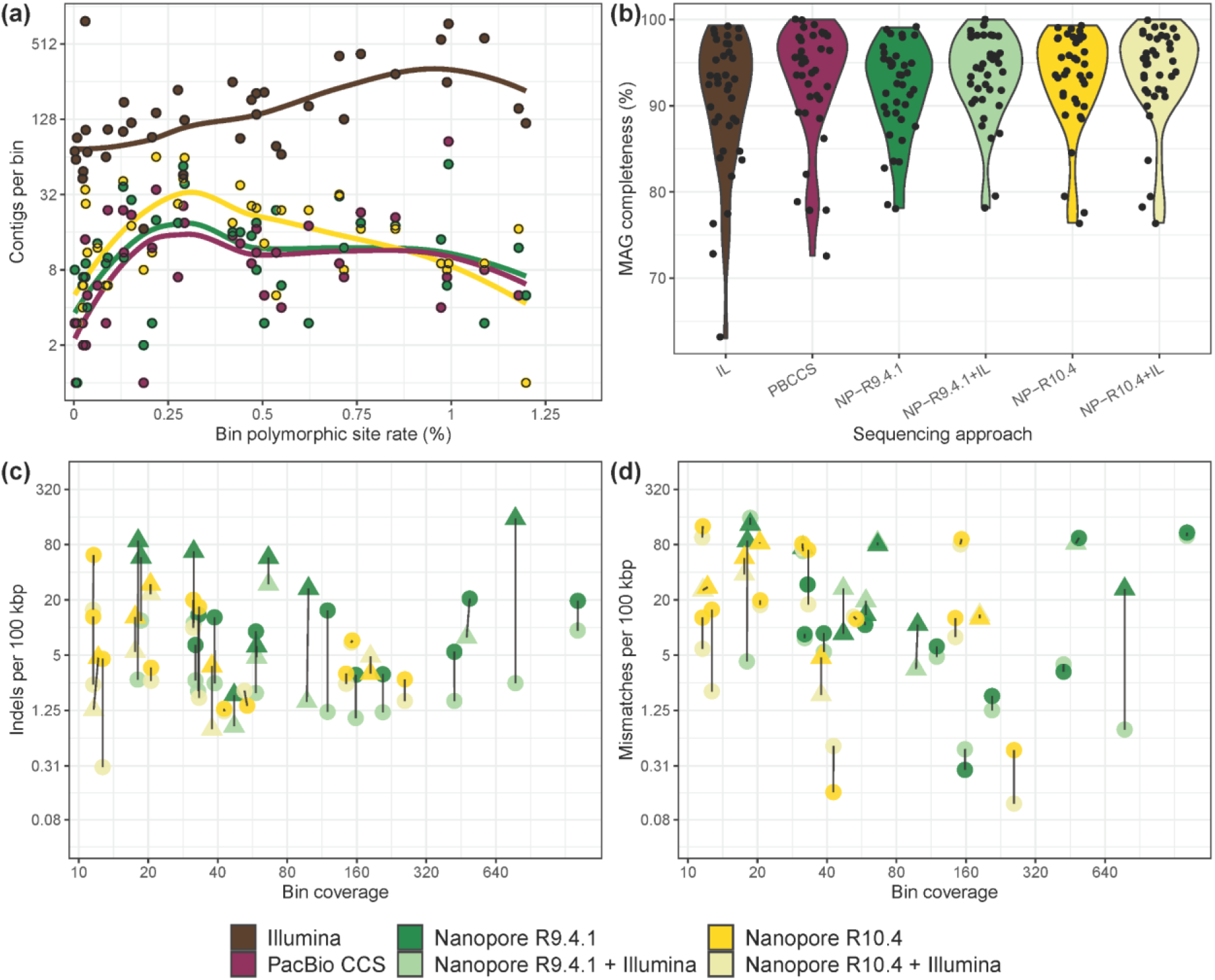
Comparison of bins from different sequencing approaches. **a)** MAG fragmentation (log-scaled) at different bin SNP rates in PacBio CCS MAGs. **b)** Genome bin completeness estimates for different sequencing platforms. IL — Illumina, NP — Nanopore, PBCCS — PacBio CCS. Bin **c)** indel and **d)** mismatch rates (log-scaled) for MAGs from Nanopore sequencing with and without Illumina read polishing, compared to MAGs from PacBio CCS. The presented bin coverage on the x axis (log-scaled) is for the corresponding Nanopore chemistry type. HQ MAGs are represented by circle, while triangles denote MQ MAGs. For all figures, only the bins that were clustered together between all the different sequencing platforms (see Materials and methods) are presented.

**Figure S7:**
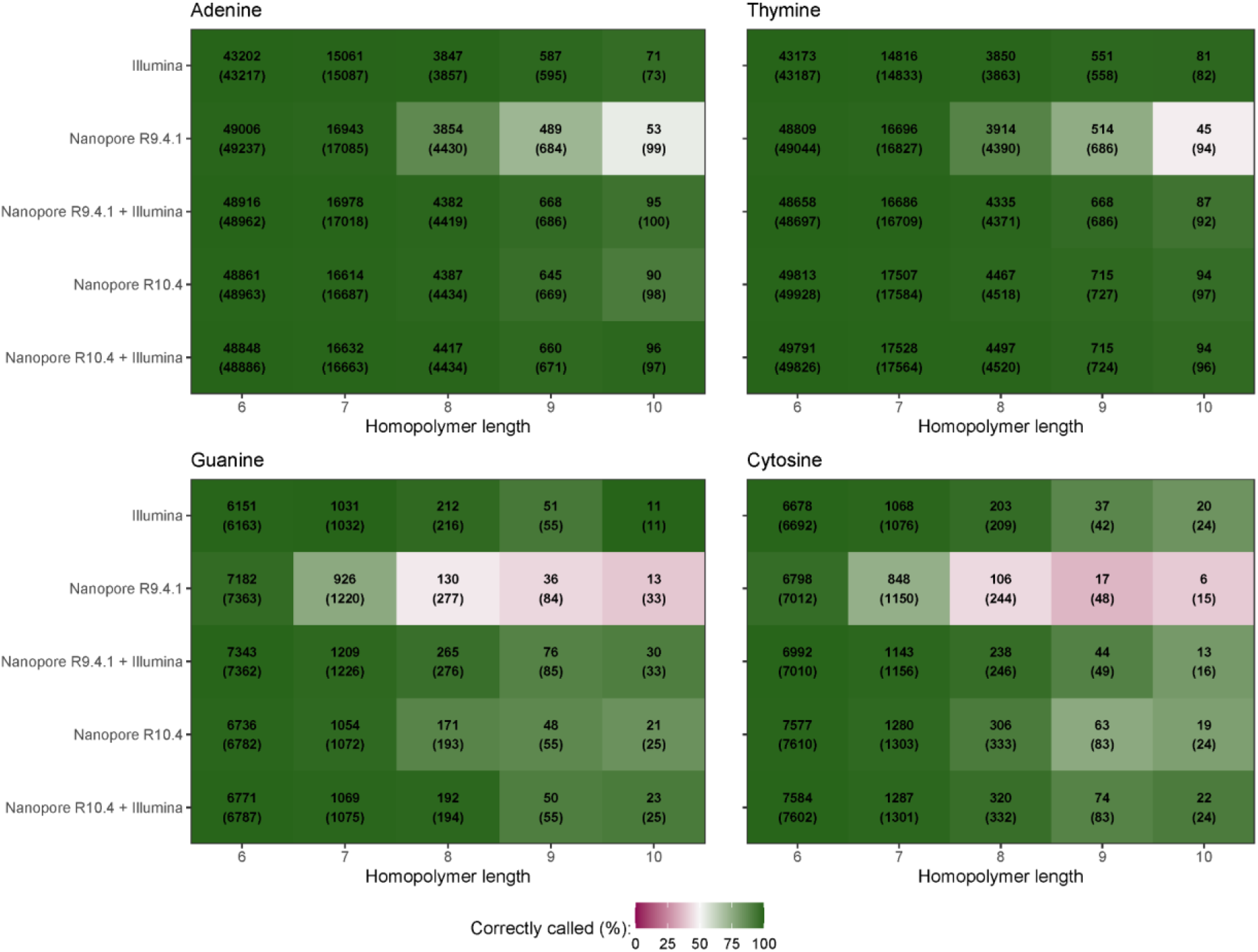
Homopolymer calling estimates in metagenomes (consensus sequences) from different sequencing platforms. Values in the heatmap show observed homopolymer counts estimated to be called correctly at a given sequence length. The total count of homopolymers (called correctly and incorrectly) are in brackets. Only the contigs for bins that were clustered together between different platforms were used to generate values for the plot.

**Table S1:** Sequence statistics for the Zymo HMW Mock using different sequencing platforms. Estimated modal read accuracy is measured using the reported Q-score for each read type. Observed modal read accuracy was measured by read-mapping to the reference genomes.

|  | **Illumina** | **Nanopore**<br>**R9.4.1** | **Nanopore**<br>**R10.4** |
| --- | --- | --- | --- |
| **Total read count** | 48,123,500 | 8,846,993 | 22,452,567 |
| **Total yield (Gbp)** | 7,2 | 31,6 | 52,3 |
| **N50 (bp)** | 151 | 14,018 | 5,992 |
| **Estimated modal read**<br>**accuracy (%)** | 99.99 | 96.89 | 98.22 |
| **Observed modal read**<br>**accuracy (%)** | 99.98 | 97.59 | 99.07 |

**Table S2:** Overview of read datasets used in the study.

| Read dataset | Instrument | Yield (Gb) | Read N50 (kb) | Read count | ENA sample ID | LOT# |
| --- | --- | --- | --- | --- | --- | --- |
| IL-201104 | Illumina HiSeq | 6.2 | 0.15 | 42,727,130 | ERS7673063 |  |
| IL-201112 | Illumina HiSeq | 11.4 | 0.15 | 79,619,634 | ERS7673064 |  |
| IL-201301 | Illumina HiSeq | 7.5 | 0.25 | 31,702,618 | ERS7673065 |  |
| IL-201308 | Illumina HiSeq | 6.7 | 0.25 | 28,067,586 | ERS7673066 |  |
| IL-201502 | Illumina HiSeq | 5.3 | 0.25 | 22,351,578 | ERS7673067 |  |
| IL-201702 | Illumina HiSeq | 15.9 | 0.25 | 66,225,442 | ERS7673068 |  |
| IL-201705 | Illumina HiSeq | 4.9 | 0.25 | 20,492,240 | ERS7673069 |  |
| IL-201707 | Illumina HiSeq | 5.5 | 0.25 | 23,663,146 | ERS7673070 |  |
| IL-201804 | Illumina MiSeq | 3.2 | 0.3 | 11,981,252 | ERS7673071 |  |
| IL-202001 | Illumina MiSeq | 13.3 | 0.3 | 47,091,904 | ERS7673072 |  |
| PB-202001 | PacBio Sequel II | 15.3 | 15.4 | 992,914 | ERS7673073 |  |
| R9-202001 | MinION Mk1B | 35.2 | 5.9 | 10,266,261 | ERS7673074 |  |
| R10-202001 | MinION Mk1B | 13.0 | 6.4 | 3,646,771 | ERS7673075 |  |
| R104-202001 | GridION | 14.0 | 7.5 | 3,514,955 | ERS7672969 |  |
| IL-ZYMO | Illumina MiSeq | 7.5 | 0.15 | 49,774,986 | ERS8296812 | ZRC195845 |
| R941-ZYMO | MinION Mk1B | 32.0 | 1.8 | 8,851,918 | ERS8296813 | ZRC195845 |
| R104-ZYMO | PromethION | 5.2 | 7.5 | 18,831,686 | ERS8296814 |  |

**Table S3:** CMSeq SNP calling statistics for the Zymo mock reference sequences.

|  | **Covered**<br>**bases (Mb)** | **Polymorphic**<br>**bases (bp)** | **Polymorphic**<br>**rate** |
| --- | --- | --- | --- |
| **Bacillus subtilis** | 4.0 | 10 | 2.5e-06 |
| **Enterococcus faecalis** | 2.8 | 113 | 4.0e-05 |
| **Escherichia coli** | 4.8 | 1156 | 2.4e-04 |
| **Listeria monocytogenes** | 3.0 | 80 | 2.7e-05 |
| **Pseudomonas aeruginosa** | 6.8 | 1222 | 1.8e-04 |
| **Salmonella enterica** | 4.8 | 41 | 8.6e-06 |
| **Staphylococcus aureus** | 2.7 | 18 | 6.6e-06 |

